# A Computational Model of Action Specification in the Basal Ganglia

**DOI:** 10.1101/2025.08.12.669938

**Authors:** Madeleine Bartlett, P. Michael Furlong, Terrence C. Stewart, Jeff Orchard

## Abstract

The basal ganglia has traditionally been modelled as a system that represents the available, discrete action space using a finite set of distinct and non-overlapping representational units. This limits these models from addressing questions around how the basal ganglia is involved in action specification: the selection of continuously valued action dynamics such as speed and vigor. In this article we present a novel computational model of the basal ganglia which incorporates vector-symbolic algebras in order to represent continuous action spaces. The stages of model development are presented along with simulation experiments to test the basic properties of the model. This work represents a promising foundational step in providing a mechanistic account for neuroscientific and behavioural evidence implicating the basal ganglia in action specification, thereby filling a gap in the literature.

## 1 Introduction

Models of action selection in the basal ganglia have, to-date, mostly modelled selection using relatively simplistic neural representations (e.g. [1–5]). In particular, models formulate selection as a process between categorically distinct actions represented using non-overlapping representational units. This, however, has limited these models from accounting for evidence suggesting that the basal ganglia also plays a role in the *specification* of action kinematics, which involves selection from continuous action spaces (such as speed [6–9], and vigor [10–12]). In this work we develop and test a novel model of the basal ganglia capable of selecting from both continuous and discrete action spaces, a role which we also refer to as *action specification*. We leverage techniques from Vector Symbolic Algebras (VSAs), also known as Hyper-Dimensional Computing, to allow us to encode distributions over continuous variables using vectors that can be represented by populations of spiking neurons. The resulting model provides a biologically plausible first look at how the basal ganglia might achieve action selection in continuous action spaces.

The basal ganglia is a collection of nuclei in the midbrain consisting of the striatum, substantia nigra pars compacta (SNc), subthalamic nucleus (STN), globus pallidus externus (GPe), globus pallidus internus (GPi), and substantia nigra pars reticulata (SNr) (see Figure 1). It is a site of special interest to behavioural and neuroscience researchers, due to findings that have implicated the basal ganglia in a wide array of behaviours including action selection [13–16], learning [17–20], decision-making [21, 22], and language [23–26]. The basal ganglia has also been implicated in the pathophysiology of several psychiatric and neuro-degenerative disorders including Parkinson’s Disease [27–29], Tourette’s Syndrome [30–32], Major Depressive Disorder [33–35], and Attention-Deficit/Hyperactivity Disorder [36, 37]. Models of basal ganglia function are, therefore, of great importance for providing insight into the development of these disorders, as well as their interventions. Continuing to push for higher fidelity in models of the basal ganglia is imperative for avoiding the drawing of incorrect or incomplete inferences.

**Fig. 1.**
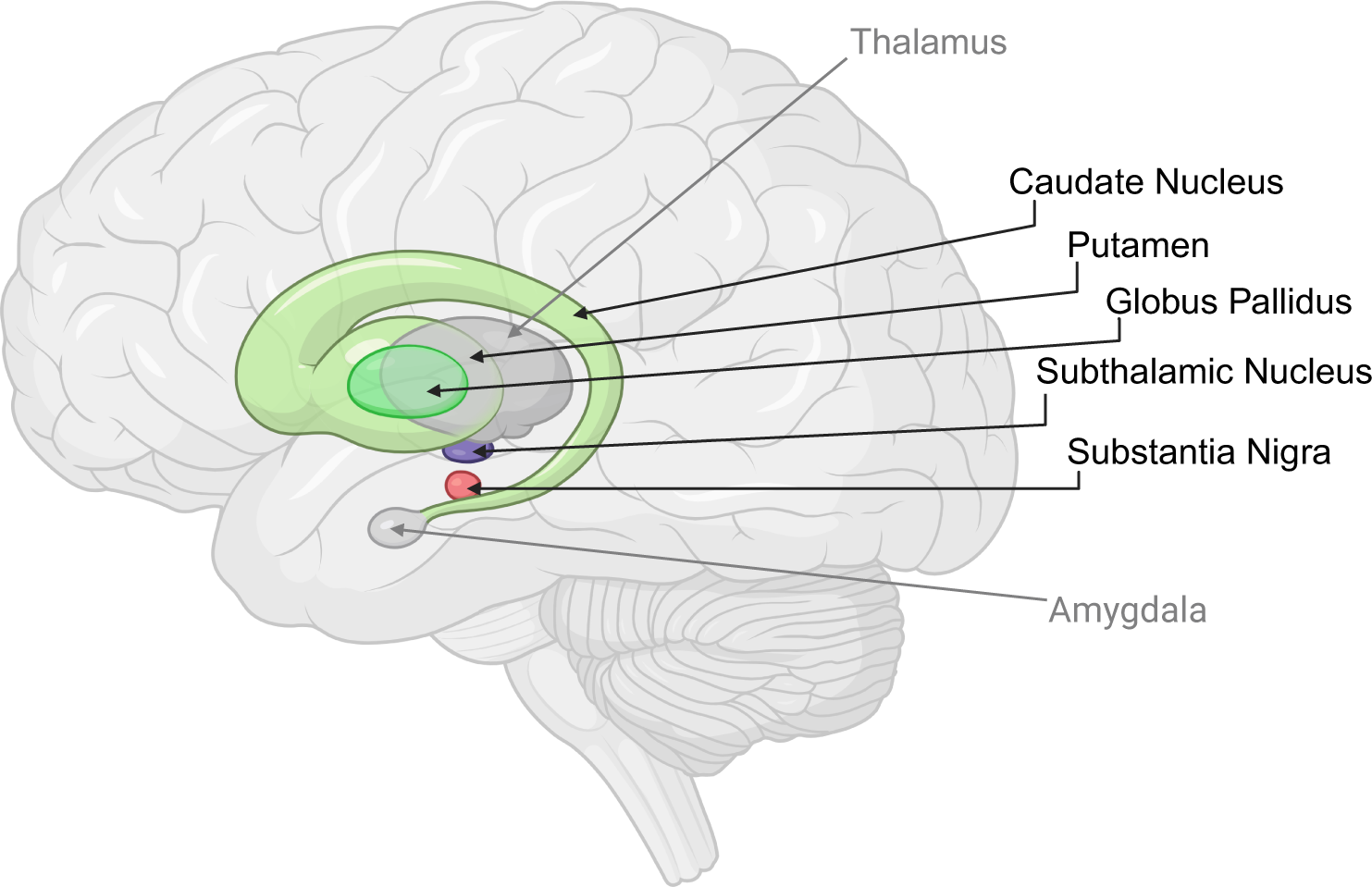
Created in BioRender. Bartlett, M. (2025) https://BioRender.com/0fus9m9. Lateral view of the brain showing the structures that make up the basal ganglia (caudate nucleus, putamen, globus pallidus, subthalamic nucleus and substantia nigra) and some related structures (thalamus and amygdala).

The majority of models to-date rely on localist encodings of necessarily discrete action spaces. Let’s first define what is meant by ‘localist’ vs ‘distributed’ encodings, and the difference between ‘discrete’ and ‘continuous’ action spaces. In the context of models of action selection in the basal ganglia, a localist encoding refers to a relatively stringent, one-hot style for representing actions. Each action is represented by a unique representational unit. These representational units are non-overlapping such that each unit is only involved in representing one action. In an action selection system that uses this representation, when changing from selecting one action to another only two units change their activity, one turning on and one turning off. The highly influential model presented by Gurney et al. [2, 38] (hereafter: the Gurney-Prescott-Redgrave (GPR) model) adopts localist encodings in the form of distinct, non-overlapping channels of neurons for representing each available action. Similarly, models developed by Berns and Sejnowski [1], Frank [4], and Lindahl and Kotaleski [5] all adopt a localist method for encoding actions. An important, limiting factor for these models is that it is necessary to introduce additional units in order to add more actions into the action repertoire. In fact, it is necessary to introduce as many new units as actions to be represented.

In contrast, distributed encoding involves overlapping groups of units for representing different actions. That is, one unit might be active in representing multiple actions, but the collection of active units will differ from action to action. As the action to-be-represented changes, multiple representational units may change their activity. A particular strength of distributed encoding is that it is capable of representing more actions than there are representational units. Thus representing more actions doesn’t necessarily require the introduction of additional units.

When considering the types of information each encoding method can represent, localist encodings can represent discrete variables very readily. Some styles of localist encoding can be used to *approximate* continuous variables by effectively discretising the continuous space. However, identifying the appropriate resolution for discretising a continuous space can be difficult and task-dependent. Furthermore, having enough localist encoding units to sufficiently cover the continuous space can be demanding of computational resources. Distributed encoding, on the other hand, is much better suited to representing continuous variables since the representational capacity is so much greater. Distributed encoding methods are also equipped to handle discrete variables and so offer a unified approach that may still reflect many of the neurological findings of action specificity in basal ganglia neuron responses.

The use of localist encoding in models of the basal ganglia is supported by findings suggesting that spatially distinct clusters of neurons are responsible for encoding semantically (or physically) separable actions. For example, the foundational work of DeLong et al. [15] in rhesus monkeys indicated that distinct populations of neurons within the striatum, STN and GPi were differentially responsible for representing movements of the forelimb, hindlimb and orofacial regions of the body. Such findings have been replicated and corroborated in primates [39, 40], humans [41, 42], and rodents [43, 44]. More recently, distinct clusters of neurons in the basal ganglia have been identified for selecting specific actions. Experimental work by Lee et al. [45] showed that stimulation of neurons in the dorsolateral striatum evoked turning behaviour in rats, whilst stimulation of neurons in the ventrolateral striatum evoked licking behaviour. Equally, Klaus et al. [46] identified specific clusters of neurons in the striatums of mice performing natural actions that activated for specific actions. In these same experiments, co-active neurons tended to be closer to one another in space. Together these findings are indicative of a local encoding of actions in striatum. As such, these localist models of basal ganglia have been successful in capturing a multitude of behaviours and have led to great insights, such as elucidating the mechanisms through which selection and decision making may be achieved [47], the effects of dopamine depletion [38], and the roles of different nuclei of the basal ganglia [48].

However, there is also evidence suggesting that the basal ganglia plays a role in action specification, or selecting the *features* of actions, or action kinematics, which are often continuously valued (e.g. speed/vigor). Sales-Carbonell et al. [49] recorded the activities of striatal neurons as mice performed a run-and-stop task. They found prolonged modulation of neuron activities while the mice were running, rather than just at the initiation and termination of running. Additionally, they identified that, at the population level, neural activity changed smoothly with the changing dynamics of the run-and-stop task, indicating that different neurons were tuned to different phases of the task. This suggests that the striatal neurons were monitoring the moment-to-moment dynamics of the animals’ movements. These results have been corroborated by others, including Fobbs et al. [6] who conducted a similar experiment but had mice perform spontaneous exploration of an open field arena. They identified neurons in the striatum that exhibited speed-tuning curves. Barbera et al. [7] identified spatially distinct clusters of neurons in the striatum of mice that elicited differential responses depending on the mouse’s locomotive speed. Thus this research suggests that the speed of an action can be decoded from the pattern of activity across the population. Indeed, Dhawale et al. [9] successfully trained a multilayer neural network to predict the velocity of head and forelimb movements based on recordings from striatal neurons.

Collectively, this work has led to the “continuous encoding model” [6] of the basal ganglia, which states that the continuous-valued features of actions – such as speed – are represented in the basal ganglia [6, 10, 49–51]. Importantly, the continuous encoding model does not preclude the idea of action-specific channels of neurons within the basal ganglia. Rather, the two representations may co-exist. For example, Klaus et al. [45] identified that, whilst distinct clusters dedicated to specific actions were evident in the striatum, actions were not necessarily encoded by discrete, non-overlapping groups of striatal neurons; similar actions were represented by overlapping sub-populations of neurons. The authors quantified the similarity between different actions of freely-moving mice by characterising the actions in terms of bodily acceleration, gravitational acceleration and rotation angle. They also quantified the similarity between striatal activity patterns for different behavioural clusters as Euclidean similarity. With these two measures, they were able to characterise the relationship between behavioural similarity and neuronal activity similarity. They observed a strong positive correlation between behavioural and neuronal similarity, indicating that there is a relationship between the degree of overlap in the neuronal representation of a pair of actions, and the behavioural similarity of those actions, characterised by their continuous, kinematic features. Similar work suggesting a hybrid localist-distributed encoding of actions and their kinematics in the striatum comes from Park et al. [12] who trained mice to perform a reach-to-pull task. Park et al. [12] manipulated the reach angle and the amount of force required to pull a joystick. They found that, whilst overlapping ensembles of neurons in the striatum were active during all reach-to-pull actions, they were able to dissociate the action features (angle and force) based on the pattern of activation within those ensembles. Together, these findings demonstrate that there may be distributed representations within the basal ganglia relating, in some way, to kinematic or continuous features of actions such as acceleration or angular rotation.

In the proposed model, we test an approach to achieving distributed representations of continuous actions in a computational model of the basal ganglia. We adopt a distributed representation of continuous action spaces wherein actions are encoded using methods from hyper-dimensional computing, aka VSA. Using VSA-based encoding approaches, we can encode items using high-dimensional vectors [52]. There are a number of advantages of this approach which we leverage for this work. First, many of the Holographic Reduced Representations (HRRs) and Fourier HRRS (FHRRs), preserve similarity relationships such that items that are similar in the domain space will be nearby to one another in the vector space [52–54]. Thus, when we encode a range of speeds, for example, as our continuous action space, the similarity between different speeds will be preserved in the vector representation. A second advantage we leverage is the VSA *bundling* method, which combines a group of vectors together to represent the set, often via some form of vector addition. Bundles in VSAs have been shown to have relationships to probability distributions [55, 56]. The operations used to construct bundles are, importantly, dimension-preserving, meaning that adding new items to a bundle does not *necessarily* require additional computational resources to represent or store (although bundling can result in noise that might reduce the clarity of the representation). Using bundling, we can construct objects that represent distributions of salience (i.e. the likelihood of an action being selected) over an action space. A final advantage we leverage is that, with some VSAs, it is possible to encode continuous variables. Different VSAs leverage different encoding and binding methods for encoding continuous variables. Encoding of continuous variables is done using circular convolution in HRRs, or equivalently, element-wise multiplication in FHRRs (which use complex-valued vectors) [57].

In this work, we adopt HRR as the chosen VSA, which has been used as a computational model in the Semantic Pointer Architecture (SPA) [58]. Within SPA, symbol-like information is represented in high-dimensional vectors referred to as Spatial Semantic

Pointers (SSPs). The bundle vectors generated via this approach can be represented in a distributed fashion by populations of neurons whereby each neuron’s activity is a weighted sum of the bundle vector (Figure 2). This is further illustrated in Figure 3 which depicts four salience distributions, encoded as bundle vectors, and then represented by an ensemble of 100 leaky integrate-and-fire neurons. The rate plots illustrate that each distribution is represented by overlapping groups of active neurons within the population. Thus this approach suits our needs, encoding distributions of salience over continuous (or discrete) action spaces as objects that can be represented by a single population of neurons. Furthermore, as shown in Figures 2 and 3, the neural representation can be made to be distributed in nature, rather than localist, allowing us to account for the findings of Klaus et al. [46] and Park et al. [12] discussed above. We follow the method introduced in [59] for encoding the salience-action distribution as a vector or SSP. In constructing the vector bundle, we start with a continuum of actions in the set A = [a_min_, a_max_]. Let a*_i_*, i = 1, …, n, be n regularly spaced samples in A and let ϕ(a*_i_*) be the SSP representing the value a*_i_*. If each action a*_i_* is associated with a salience s*_i_*, then we can weight each ϕ(a*_i_*) by its associated salience s*_i_* to give s*_i_*ϕ(a*_i_*). We then use the HRR bundling operation, superposition, to bundle the weighted vectors together into a single vector 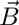 such that

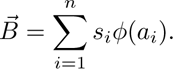

**Fig. 2.**
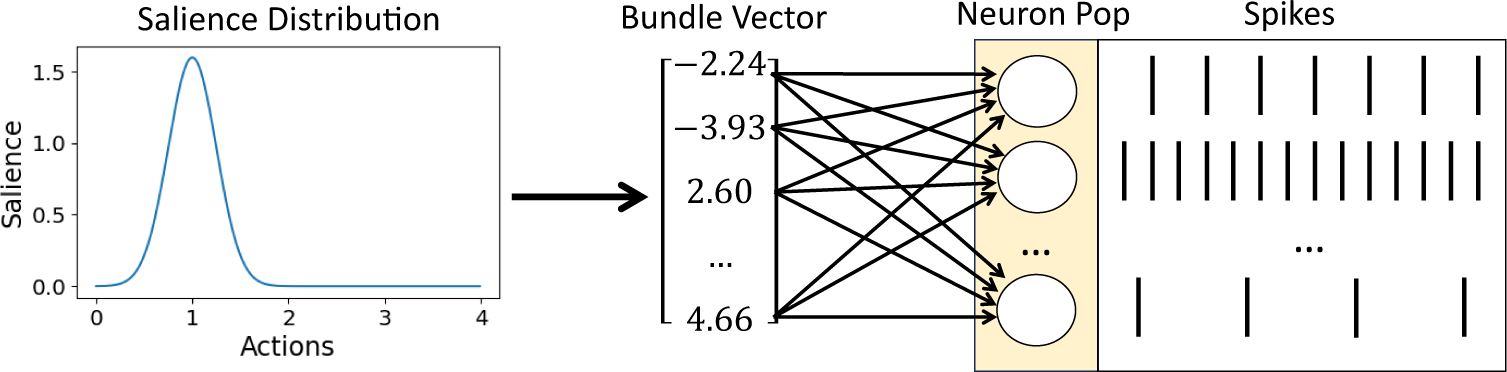
Illustration of the process of encoding a salience distribution over a continuous action space. From left to right; distribution encoded as a bundle vector. All-to-all connections (with random encoding weights) from each dimension of the vector to the neurons in a population of spiking neurons. The activity of each neuron in the population representing the bundle vector. We plot the neuron activities here as spikes for clarity, but in simulation we use rate-approximation for simplicity.

**Fig. 3.**
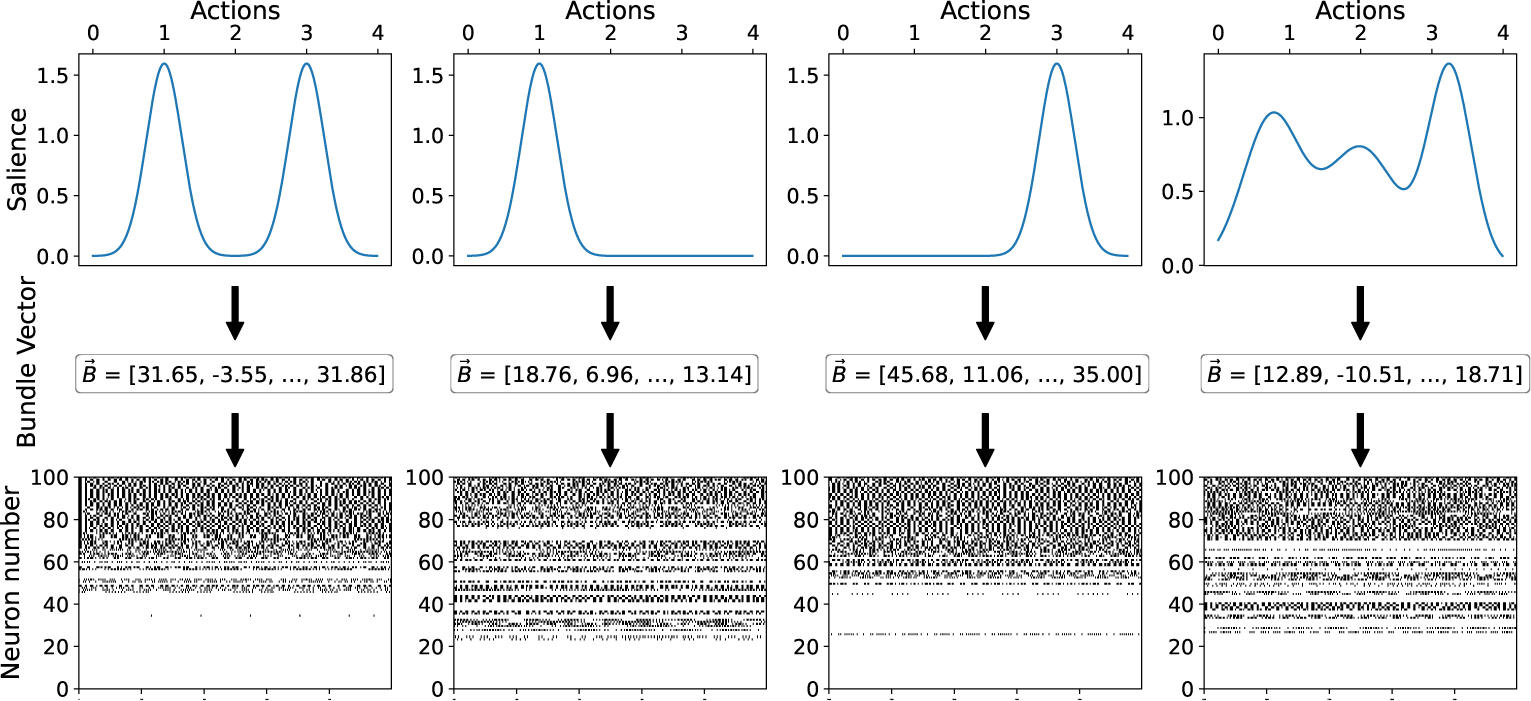
Illustration of the process of encoding four different salience distributions over a continuous action space. The top row shows the salience distributions. The second row shows part of the 512D SSP vector embedding of the distribution. The bottom row shows the neuron firing rates of an ensemble of 100 Leaky Integrate-and-Fire Rate neurons representing the vector *B*. To illustrate that this results in a distributed representation of the input signal, we note that the spiking activity in column one is *not* a combination of the activity in the second and third columns where the neurons are representing each half of the bimodal distribution.

With this formulation, it is non-trivial to have the basal ganglia’s output be a single selected action, as it has been in previous models (e.g. [2, 4, 5]). The ‘selected’ action is represented as the action with the highest salience, or the location of the peak of the salience distribution. However, finding the max of a distribution is not straight-forward for neural networks. We therefore hypothesise that the role of the basal ganglia might not be to isolate a single action for selection, but rather to act as a clean-up system. Specifically, we propose that the basal ganglia’s role could be to concentrate the distribution of salience around a point and thereby limit the range of actions from which a down-stream network can select. We quantify this in Section 3.1 as both the difference in entropy before and after 2, and the difference in location of the peak before and after 1 – because the basal ganglia should not change which point in the action space has the highest salience.

Furthermore, previous models, such as the GPR, have exhibited a tendency for simultaneous selection when two action channels have the same, sufficiently-high salience [38]. This is somewhat counter to the believed role of the basal ganglia which is to prevent the simultaneous selection of incompatible actions (e.g. attempting to simultaneously wave and write with the same hand). We therefore further specify that the basal ganglia’s role should include resolving competition between equally salient actions by concentrating the salience distribution under one mode and not the other. We conduct simulation experiments to determine whether this property is held by the current model.

A final property we examine is the behaviour of the network under conditions of dopaminergic modulation. One of the roles played by dopamine during learning is to control the explore/exploit trade-off during learning by modulating the behaviour of the basal ganglia. We therefore attempt to replicate the experiments in Humphries et al. [60]. In this prior work, Humphries et al. [60] examined whether increasing levels of dopamine in the GPR model led to more exploitative behaviours (higher probabilities of selecting the high salience action) compared to when dopamine levels were low. We replicate this experiment with our novel model in order to determine whether this property holds.

The first step in constructing the proposed model of the basal ganglia is to identify the network dynamics that can achieve this entropy-reducing operation. We conducted an initial optimisation experiment wherein we identified several candidate networks for concentrating the salience distribution and compared how well then did at reducing the entropy of the distribution, whilst maintaining the same peak location. We identify that the Dynamic Neural Field (DNF) network achieved the best performance. We therefore incorporate the DNF into a model of the basal ganglia and, in a set of simulation experiments where we test a number of model properties to provide preliminary validation of the model. This work constitutes a major foundational step in modelling the role of the basal ganglia in specifying continuously-valued action kinematics.

## 2 Methods

### 2.1 Experiment 1

#### 2.1.1 Action-Place Cells

In many of the networks compared for entropy-reduction it was necessary to decode the action-salience bundle in order to act on the salience values separately of the vectors encoding the actions. We achieved this decoding by using a population of neurons with place-cell like activities. We refer to these neurons as action-place cells. Biological place cells are neurons found in the hippocampus that become active when the agent enters a particular location in their environment. Using VSA techniques, one can create a population of place cells by sampling locations from across an environment, encoding those locations as high dimensional vectors (e.g. SSP), and using those vectors as the synaptic weights for a population of neurons such that each neuron’s receptive field is centred around the corresponding location within the data space or environment. Because VSAs, preserve similarity in the domain space, these neurons will display some activity whenever the agent enters a location that is nearby to the location matching their preferred stimuli. Each neuron will fire most strongly for the specific location used to define its encoder.

To create the action-place cells, we sampled values from the continuous action domain, and used those values, encoded as SSPs, as the encoders for a population of neurons. As a result, each neuron’s receptive field was tuned to a single SSP encoding a single action. The effect of this is a population of neurons whose activities reflect the dot product of the matrix of encoded SSP values and the input signal. When this input signal is the SSP bundle used to encode the salience distribution over a continuous action space, the result of this dot-product is a vector containing the salience of each of the decoded actions. Thus, the activity of this neuron population represents the distribution of saliences across the action space.

Once decoded, we can perform transformations on the salience distribution, such as lateral inhibition, and pass the new distribution through the transpose of this encoder matrix in order to re-encode the salience distribution as an SSP bundle. Due to the small distances between encoded actions, and the fact that the similarity curves for each SSP overlapped with up to 50 of its neighbours on either side, we do not consider this a localist encoding. This is because, despite the bundle being constructed via a weighted sum of a *finite* set of vector-encoded actions, it is possible to query the bundle to find out the height of the distribution at points in-between those vectors. For example, if we construct the bundle

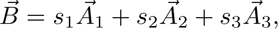

we can find the value of s_1.5_ by calculating the dot product between the bundle 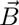 and the vector 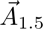,

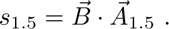

Thus, loss of a neuron does not necessarily result in an inability to alter the salience or select the action encoded by that neuron. In the following networks, the action-place cells consisted of a population of 400 LIF-Rate neurons with encoders set to a set of 400 SSPs encoding actions sampled at evenly spaced intervals across the action space.

#### 2.1.2 Networks

Networks were constructed in Python using the Neural Engineering Framework (NEF) and Nengo library [61]. Schematics for each network are shown in Figure 4.

**Fig. 4.**
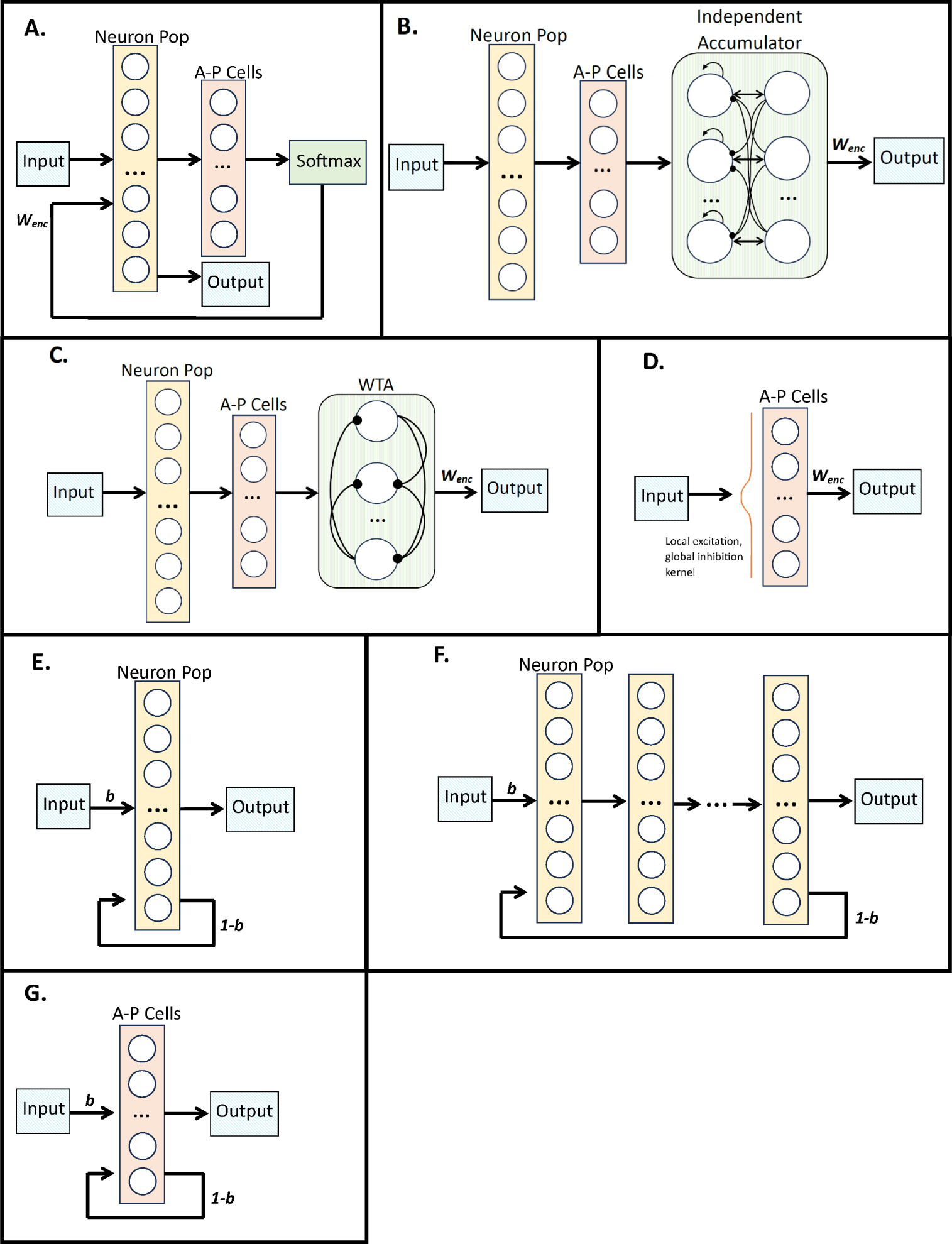
Schematic of A) Modern Hopfield network, B) Independent Accumulator network, C) Winner-Take-All network, D) Dynamic Neural Fields network, E) Shallow Attractor, F) Deep Attractor, and G) Shallow Attractor with action-place cells (AP Cells). Arrows denote excitatory connections, lines ending in circles denote inhibitory connections.

We implemented a modern Hopfield network, as defined in [62]. The states of the neurons are updated using

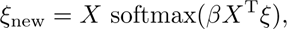

where ξ represents the network state (activities), X stores the set of target patterns to be memorized, and β is a constant defining the temperature of the softmax. The Hopfield network has two layers, one initialized to 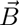, and the other set up as an array of 400 action-place (AP) cells; each AP cell corresponds to one column of X, and thus one location in the action space. The input bundle 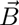, which represents a salience distribution, is sampled at each of the target locations. These values are stored in the AP cells. Then a softmax function is applied to the AP cell neuron activities in order to resolve the activity pattern to a more concentrated distribution over the action space. A second network was the Independent Accumulator (IA) network [63], also from the Nengo SPA library (Figure 4B). This network uses lateral inhibition between separate neuron populations to resolve competition. The IA is a two-layer network wherein the first layer is made up of a set of integrators, or accumulators, that accumulate the incoming signal. A second, thresholding layer applies the threshold such that once the threshold is exceeded, a feedback connection stabilises the current choice whilst inhibiting the others. For these experiments, the threshold applied to the IA = 0.8. The bundle was decoded by a population of action-place cells, and the salience distribution was then passed to the accumulator network. The output was then re-encoded as an SSP bundle.

A similar network used was the Winner-Take-All (WTA) network from the Nengo Spa library (Figure 4C). In this network, lateral inhibition is applied between a set of separate, thresholding neuron populations in order to induce competition. For each neuron population, if the inputs are below a given threshold, the population’s output will be 0, and for inputs above that threshold the population’s output will be equal to the input. For these experiments, the threshold = 0.001. The SSP bundle is decoded to the salience distribution via a population of action-place cells. The salience distribution is then passed as input to the WTA network. The output from the WTA is then re-encoded as an SSP bundle in a neural population via a weight matrix of the transpose of the action-place-cell encoders.

We also compared a DNF network, wherein a local excitation and global inhibition kernel is applied directly to the activities of a population of action-place cells (Figure 4D). The dynamic neural fields theory is described in Schoner and Spencer [64].

In order to experiment with a network that might be able to operate within the SSP space, i.e. without relying on decoding the salience distribution from the bundle via action-place cells, we included both shallow and deep attractor networks. In the first instance, we implemented a shallow attractor network consisting of an input layer, connected to a recurrently connected neuron layer (see Figure 4E). The weights on the recurrent connection were trained in order to create the attractor dynamics, such that the network would try to converge to an SSP bundle representing a more concentrated distribution of saliences.

Using the nengo deep learning library, we also constructed a deep attractor network [65, 66]. This network consisted of multiple layers, with the final layer having a recurrent connection with the first (see Figure 4F).

Finally, in order to test whether the presence of the action-place cells was an essential feature for improved performance, we also implemented a shallow attractor network where the recurrently connected layer consisted of a population of action-place cells. The SSP bundle was decoded into the salience distribution, and the attractor dynamics attempted to converge to a concentrated distribution centred at the location of maximum salience.

During training of all of the attractor networks, the networks were shown on a set of N random bundles and trained to produce a vector encoding a single action, specifically the action with the highest salience in the input distribution.

#### 2.1.3 Optimisation

Table 1 presents the hyperparameters that were optimised for each network, and the ranges of values that were tested. Using the Tree-structured Parzen Estimator (TPE) algorithm to search the parameter space, we had optuna run for a total of 100 trials per network. Each of those trials consisted of selecting the values for the hyperparameters, training the network, and then testing performance by presenting the trained network with 50 different input bundles.

**Table 1.** Ranges of hyperparameters explored for tuning the networks.

| Parameter | Description (Networks) | Ranges Explored |
| --- | --- | --- |
| Input weight | Bias weight applied to the input vector (All) | $[1e^{-9}, 2.0]$ |
| Ensemble neurons | Number of neurons in first hidden layer (All but Shallow Attractor with Action-Place cells) | 1000, 2000, 5000 |
| Temperature | Temperature value for the softmax (Hopfield) | $[0.1, 10]$ |
| Threshold | Decision threshold (Independent Accumulator, Winner-Take-All) | $[0.1, 1.5]$ |
| Intercept Width | How widely distributed the intercepts are (Independent Accumulator, Winner-Take-All) | $[1e^{-3}, 1.0]$ |
| Net neurons | Number of neurons in each layer of the internal network (Independent Accumulator, Winner-Take-All) | 50, 100, 200, 500 |
| Inhibition scale | Scaling of the lateral inhibition (Winner-Take-All) | $[0.1, 5.0]$ |
| H | Resting level of neural activity (DNF) | $[-30, 30]$ |
| Global inhibition | Strength of global inhibition applied to whole neural field (DNF) | $[0, 20]$ |
| Tau | Time constant - how quickly the neuron activities decay (DNF) | $[1e^{-5}, 0.5]$ |
| Excitation kernel | Amplitude of the excitation kernel (DNF) | $[0, 20]$ |
| Excitation width | Width of excitation kernel (DNF) | $[0, 20]$ |
| Inhibition kernel | Amplitude of the inhibition kernel (DNF) | $[0, 20]$ |
| Inhibition width | Width of inhibition kernel (DNF) | $[0, 20]$ |
| N layers | Number of hidden layers (Deep Attractor) | $[2, 10]$ |
| Learning rate | Learning rate (Deep Attractor) | $[1e^{-9}, 0.5]$ |

Performance was measured in terms of 2 metrics. The first was error, calculated as the Root Mean Square Error (RMSE) of the distance between the peak of the salience distribution used as the input, and the peak of the salience distribution produced as output by the network. This error term was used to indicate the accuracy of the network by measuring whether the action with the highest salience was the same. We calculated the error term as

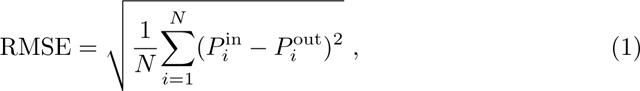

where P ^in^ is the value of the action where the peak of the input distribution occurs, and P ^out^ is the value of the action where the peak of the output distribution occurs, the subscript i indicates trial i, and N = 50. Here, values closer to 0 indicated less error and therefore better performance.

The second metric was the change in entropy, denoted ΔH. For each of the 50 test inputs, we calculated the difference between the entropy of the input distribution and the entropy of the output distribution (ΔH). We then calculated the mean ΔH using,

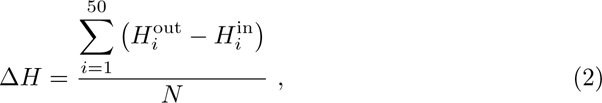

where 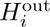 is the entropy of the i^th^ output distribution, and 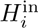 is the entropy of the i^th^ input distribution. This metric indicates the degree to which the network has concentrated the salience distribution (lowering entropy), yielding smaller, more negative ΔH values indicating better performance.

For the purposes of these experiments, the output of the basal ganglia is decoded into a salience distribution using the inner-product to calculate cosine similarity,

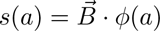

where s(a) is the salience of action a, 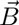 is the bundle produced by the basal ganglia model, and ϕ(a) is the vector encoding of action a. The full salience distribution is estimated by repeating this calculation for a set of actions a ∈ A. It was necessary to convert the decoded distribution before calculating entropy because entropy requires the distribution to contain only positive values, and the result of decoding using cosine similarity will contain negative values. We therefore squared the cosine similarity to make it non-negative and then normalised the result. The objective function for the optuna optimisation was to minimize both of these metrics, where the two metrics were equally weighted.

Based on the results of this optimisation study, we selected the DNF with the hyperparameter values shown in table 2 as our model of basal ganglia function for the remainder of the experiments.

**Table 2.** Best hyperparameter values identified by Optuna for the DNF network.

| Symbol | Hyperparameter | Value |
| --- | --- | --- |
| $H$ | Resting neural activity | -3.009 |
| $GI$ | Global inhibition | 8.641 |
| $\tau$ | Rate of neural activity decay | 0.047 |
| $K_e(t)$ Excitation kernel | 9.421 | |
| $\sigma(K_e(t))$ | Width of excitation kernel | 8.535 |
| $K_i(t)$ | Inhibition kernel | 1.484 |
| $\sigma(K_i(t))$ | Width of inhibition kernel | 4.748 |

### 2.2 Experiment 2

#### 2.2.1 Model

The novel basal ganglia model is presented in Figure 7. Here we adopt the same architecture as the model presented by Gurney et al. [2], and later adapted by Stewart et al. [3]. The input layer of the network is the striatum, which itself consists of 2 separate neuron populations. These 2 populations correspond to the D1 and D2 neurons found in the mammalian striatum. We hypothesise that this may be the site of the DNF action of concentrating the salience distribution. Before information from the cortex is delivered to the striatum, the bundle is concatenated together with a scalar denoting the level of tonic dopamine (DA) currently acting on the basal ganglia (in the Concatenation layer). This allows us to feed in tonic dopamine as an external and adaptable signal, enabling the online adaptation of the weights from cortex to striatum. These weights differ depending on the target population within striatum. This is to reflect to opposing influence of dopamine on the activity of D1 and D2 neurons. D1 neurons fire less strongly as dopamine levels increase, thus the weights for these connections are set in order to achieve the equation

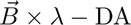

where λ is the baseline strength of the connections. In these experiments, λ = 1.0. On the other hand, the activity of D2 neurons increases as dopamine levels increase, so the weights are calculated such that the input to the D2 population is equal to

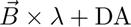

The D1 and D2 populations both consist of populations of 400 action-place cells with the DNF activation kernel applied to the neuron activities in order to concentrate the salience distribution. The salience distribution is then re-encoded as the bundle before being sent, via inhibitory connections, to the GPi (via the direct pathway) and the GPe (via the indirect pathway). The input from cortex is also sent directly to the STN, bypassing the striatum. Thus the activity of the STN represents the original, un-concentrated bundle 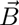. This bundle is used to encourage exploratory behaviours. By introducing a version of the distribution with a broad base, and mixing it (via excitatory connections) with the concentrated distribution, we broaden the distribution, thus providing a wider range of behaviours to the selection mechanism. We can thus change the balance between exploration and exploitation behaviours by changing the relative strengths of the indirect and direct pathways via DA.

## 3 Results

### 3.1 Experiment 1 - Hyperparameter Optimisation and Network Comparison

The first experiment set out to identify the network dynamics that are sufficient for achieving the desired entropy-reduction of the salience distribution. We compare 7 candidate networks; a modern Hopfield, an independent accumulator, a WTA, a DNF, two shallow attractors, and a deep attractor network. Schematics of the tested networks are shown in Figure 4. We use the Optuna library [67] for conducting hyperparameter optimisation.

The results of the optimisation are shown in Figure 5. In both plots, ‘best’ performance is achieved when both the RMSE and mean ΔH are minimised (i.e. the bottom left corner). The figure on the left shows the pareto-fronts calculated for each network. From this network we can see that the DNF network appears to outperform all the others.

**Fig. 5.**
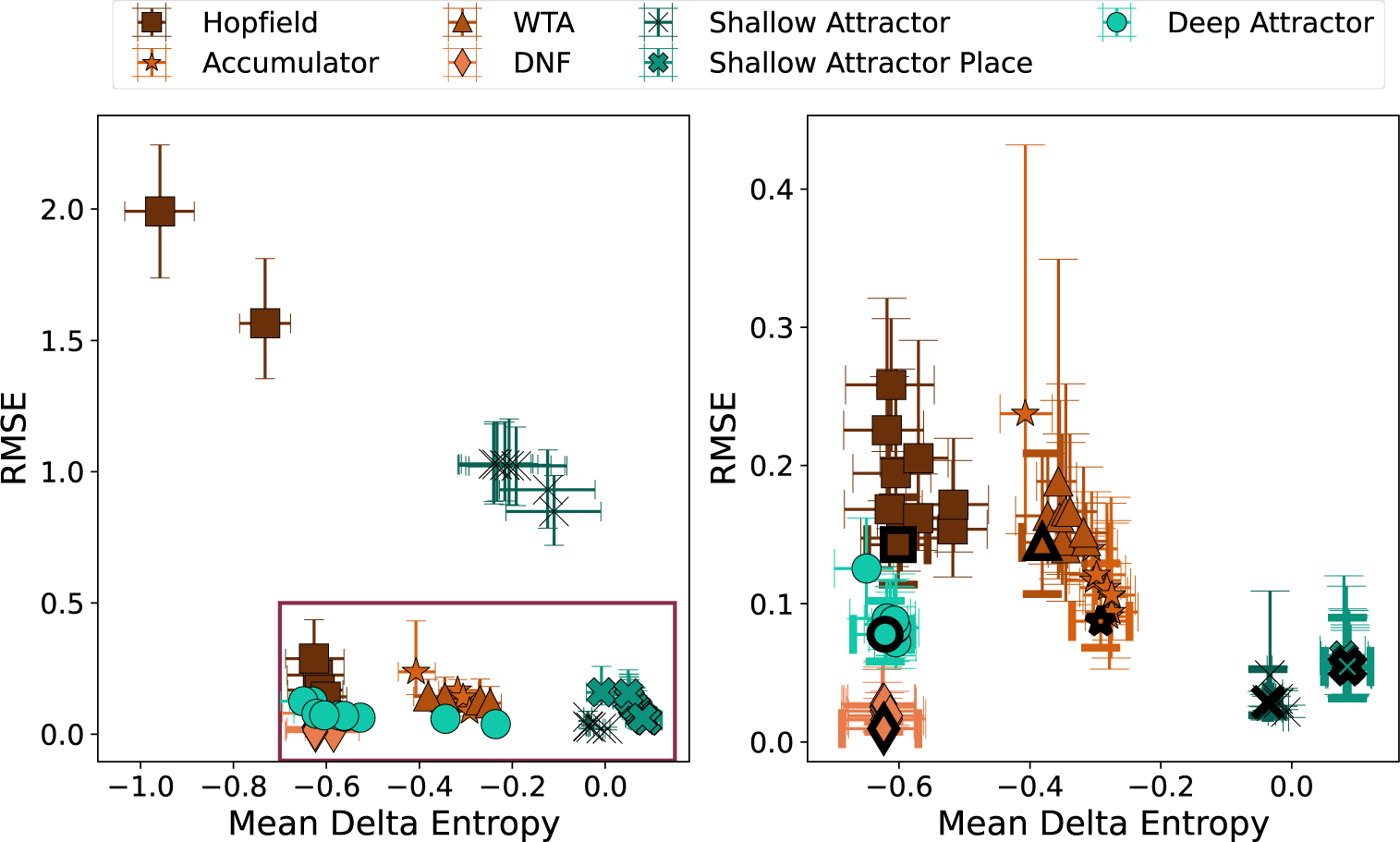
Left: Pareto-front for each network, as calculated by Optuna. Box indicates region shown in right-hand plot. Right: 10 trials that achieved the ‘best’ performance in terms of minimising both performance metrics. Error bars show 95% confidence intervals.

In order to identify which parameter set achieved the ‘best’ performance overall for each network, we calculated a single performance score,

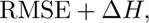

summing the two metrics together. Trials with the smallest value on this score were the best performing. The plot on the right of Figure 5 shows the top ten ‘best’ optuna trials for each network. The overall best trial is highlighted with a bolder outline around the marker. Here, we again confirm that the DNF outperformed all other networks after optimisation.

To better assess the behaviour of each network, we took the best performing parameter sets for each and tasked the networks with performing the proposed function (concentrating the salience distribution) on both a uni-modal beta distribution, and a bi-modal distribution. The results are shown in Figure 6.

**Fig. 6.**
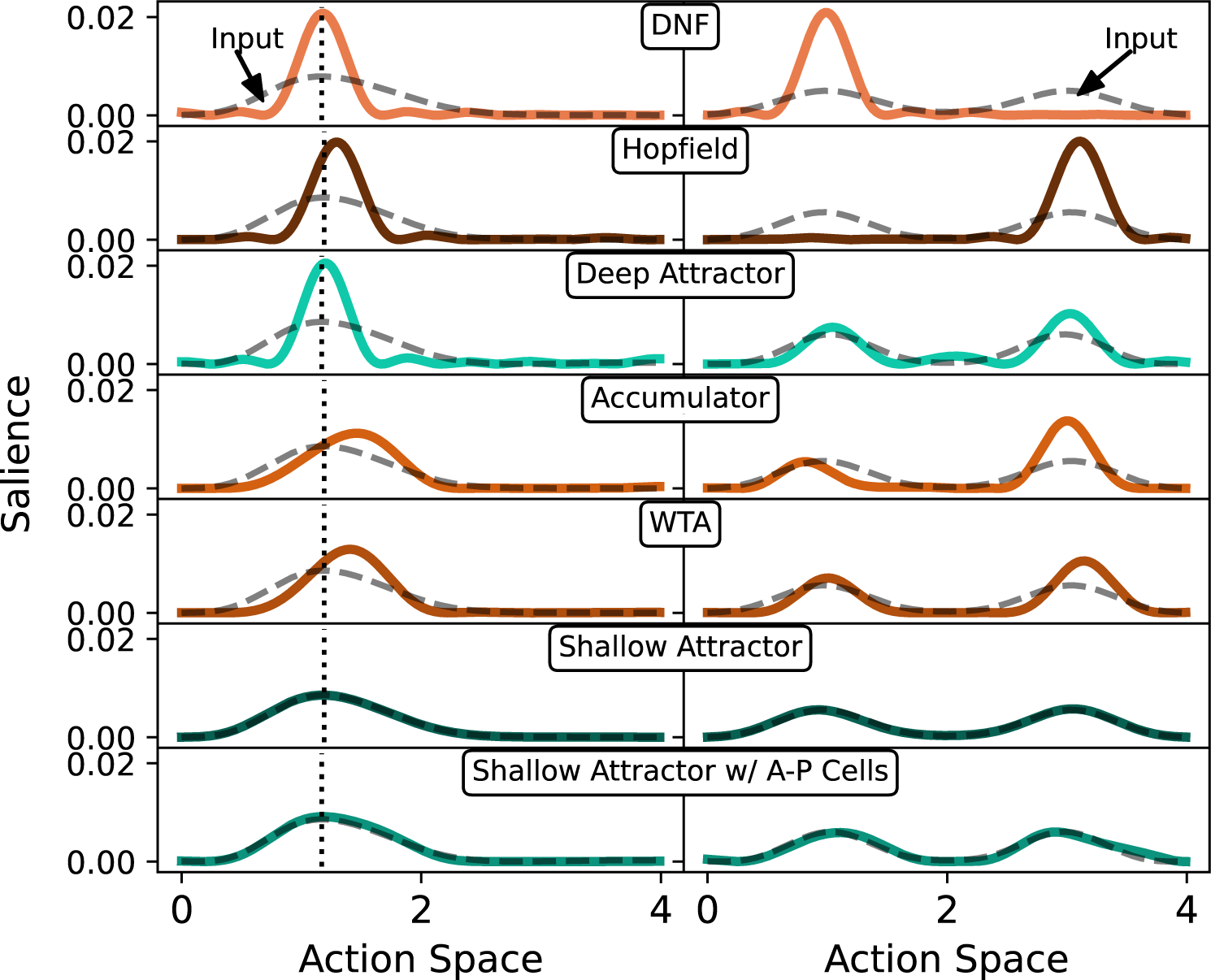
Left: Output from each network in response to uni-modal beta distribution. Right: Output from each network in response to bi-modal distribution.

Observing the behaviour of each network in response to a uni-modal distribution indicates that not only the selected DNF network, but also the hopfield and deep attractor networks, achieve the intended entropy reduction. We also examined the behaviour of each network in response to a bi-modal distribution in order to further illustrate the appropriateness of the DNF for the proposed basal ganglia model. Previous work has demonstrated that the DNF will collapse multimodal distributions to a single peak [64]. We adopt the assumption that this is desirable behaviour in a model of the basal ganglia. In a real, biological system, it is conceivable that there may be two (or more) actions with equal salience in a given context. For example, if you are approaching a pedestrian crossing on foot and there is a car approaching, your two viable options might be to slow down your walking speed to let the car pass, or to speed up and cross before the car arrives. The system must be able to choose between the two in order to avoid: inaction, attempts to perform incompatible concurrent actions (e.g. walking slowly and running at the same time), or attempts to perform the inappropriate mean action. None of the networks were exposed to bi-modal distributions prior to this test, all being evaluated and optimised on uni-modal beta distributions. As anticipated, the DNF not only performs well on the unimodal distribution, but also exhibits the desired behaviour on bimodal distributions – concentrating salience under one peak and eliminating salience under the other.

It is therefore noteworthy that the DNF and modern Hopfield networks not only perform well on concentrating the uni-modal distribution (Figure 6 Left), but are the only networks that appear to make a decisive selection between the two peaks in the bi-modal distribution. While other networks perform well on the unimodal distribution, for example the deep attractor, they fall short when it comes to the bimodal distribution, maintaining salience under both peaks.

We therefore adopted the DNF as the network dynamics for achieving action specification in the basal ganglia. The next stage of this research, then, was to incorporate the DNF into an anatomically accurate model of the basal ganglia. As part of this, we needed to identify where in the basal ganglia we believe this DNF dynamic is occurring. We then need to examine how the vector bundle is transformed by the combinations of inhibitory and excitatory connections through the basal ganglia. The next section describes the proposed basal ganglia model and findings from simulations designed to examine the properties of that model.

### 3.2 Experiment 2 - Simulation Studies

The proposed basal ganglia model adopts the same underlying architecture as the GPR model (see Figure 7). We model connections from cortex to the two populations of dopaminergic neurons in the striatum as well as to the STN. We also include the internal inhibitory and excitatory connections, and the inhibitory connections from GPi projecting to downstream nuclei (e.g. the thalamus). Rather than selecting from multiple, discrete actions, this network models a single channel encoding a distributed representation of an action-salience bundle. We employ the DNF dynamics from the first experiment in the striatum of the basal ganglia and eliminate the lateral inhibition previously implemented in the connections from STN.

**Fig. 7.**
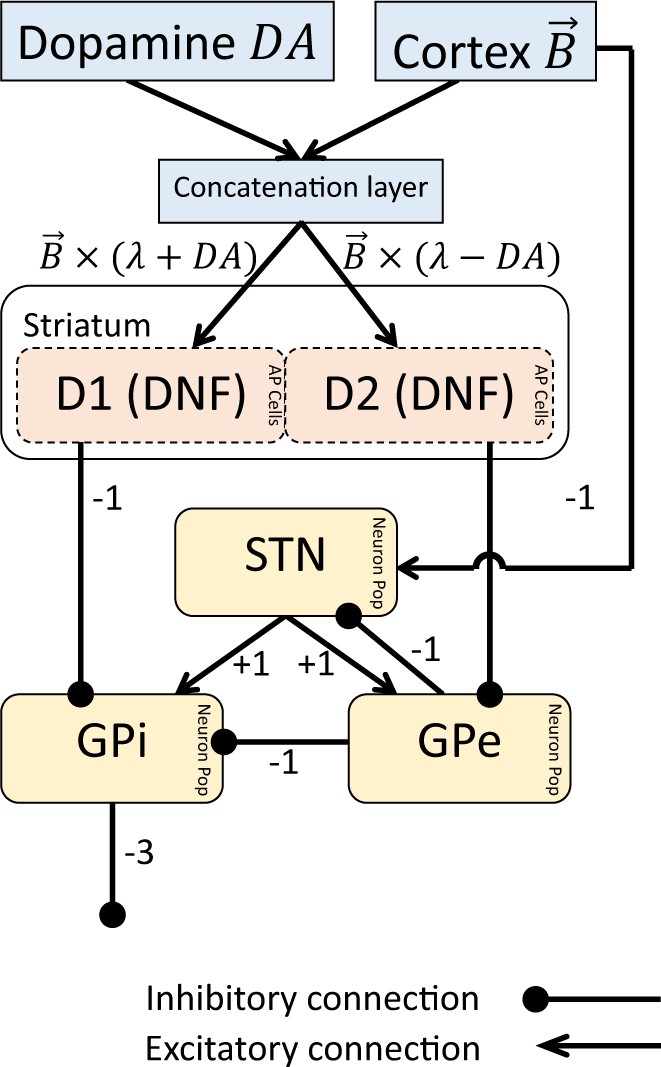
Schematic of the developed basal ganglia network incorporating the DNF network in the striatum.

To validate the model, we test some basic properties of the model.

#### 3.2.1 Property 1 - Concentrating distributions

First, we wanted to establish whether the network would successfully concentrate the distribution encoded in the vector bundle. For this experiment we generated a beta-distribution and encoded it as a bundle vector. This bundle was then used as input for the network depicted in Figure 7, where λ = 1.0 and DA = 0.0. Figure 8A shows the results of this experiment and shows that the full basal ganglia network is successful in concentrating the uni-modal input distribution.

**Fig. 8.**
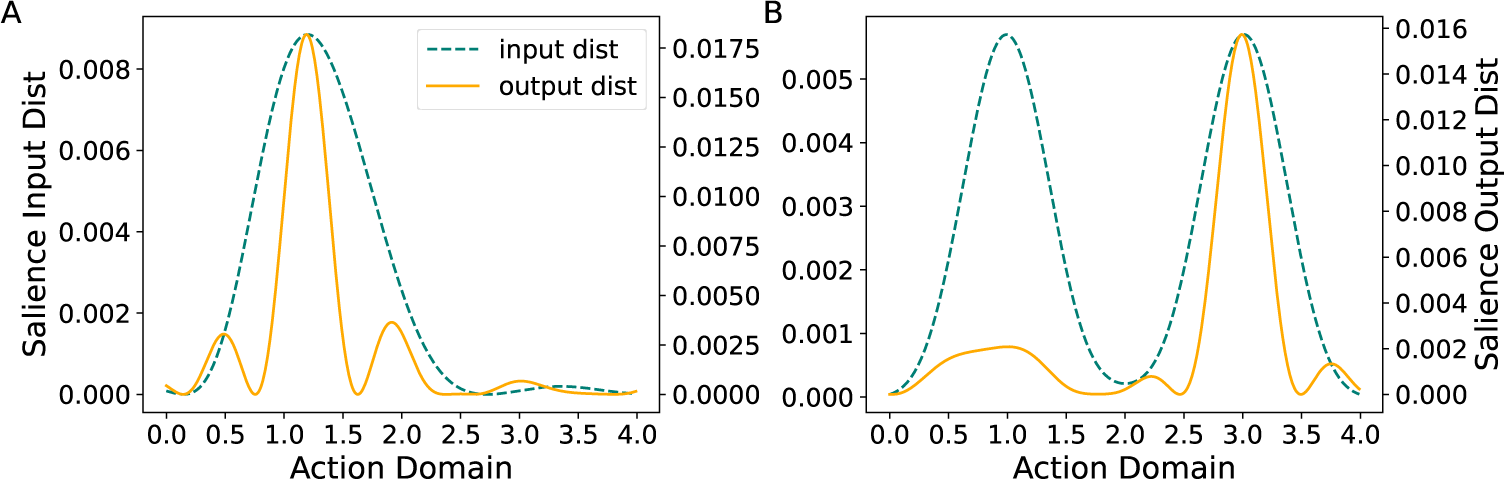
Decoded salience of basal ganglia input and output distributions when (A) input is uni-modal, and (B) input is bi-modal.

#### 3.2.2 Property #2 - Multiple options

Second, we wanted to test what the network would do with a bi-modal distribution. Figure 8B demonstrates that the basal ganglia network, when presented with an input with two equally salient regions of the continuous action space, concentrates the distribution under one of the modes. Humans and animals are often faced with scenarios where there are multiple viable options, even in the realm of continuous action features. It is important, therefore, that a model of the basal ganglia demonstrate the capability to select between these options decisively. Furthermore, we want to avoid selecting the mean of the options since that can lead to the selection of low-salience, intermediate actions.

#### 3.2.3 Property #3 - Tunable Explore/exploit Trade-off

The final property we assess is the impact of dopamine on the explore/exploit trade-off. The explore/exploit trade-off is a fundamental concept in decision-making whereby agents are required to balance the need to explore new possibilities with the need to exploit the actions that it has learned will reliably lead to goal states. It has been observed that switching from exploration to exploitation is associated with changes in tonic dopamine in the basal ganglia [60, 68]. Specifically, increased levels of dopamine are associated with an increased tendency to exploit, whilst lower levels of dopamine are associated with more exploratory behaviours. To see if our model has the same behaviour, we conducted a conceptual replication of the experiment conducted by Humphries et al. [60]. In their study, Humphries et al. [60] tested the behaviour of the GPR basal ganglia model whilst manipulating the level of dopamine in the network. They normalised and inverted the output from the network in order to interpret it as a probability distribution (see Figure 9A). Using this interpretation, they demonstrated that altering the level of dopamine in the striatum of the model had the effect of changing the balance of exploitation and exploration as represented in the probability distribution. Under conditions of low tonic dopamine, our basal ganglia model produced a relatively flat probability distribution with high entropy, whereas a higher level of tonic dopamine increased the peak of the probability distribution and reduced the entropy (Figure 9A).

**Fig. 9.**
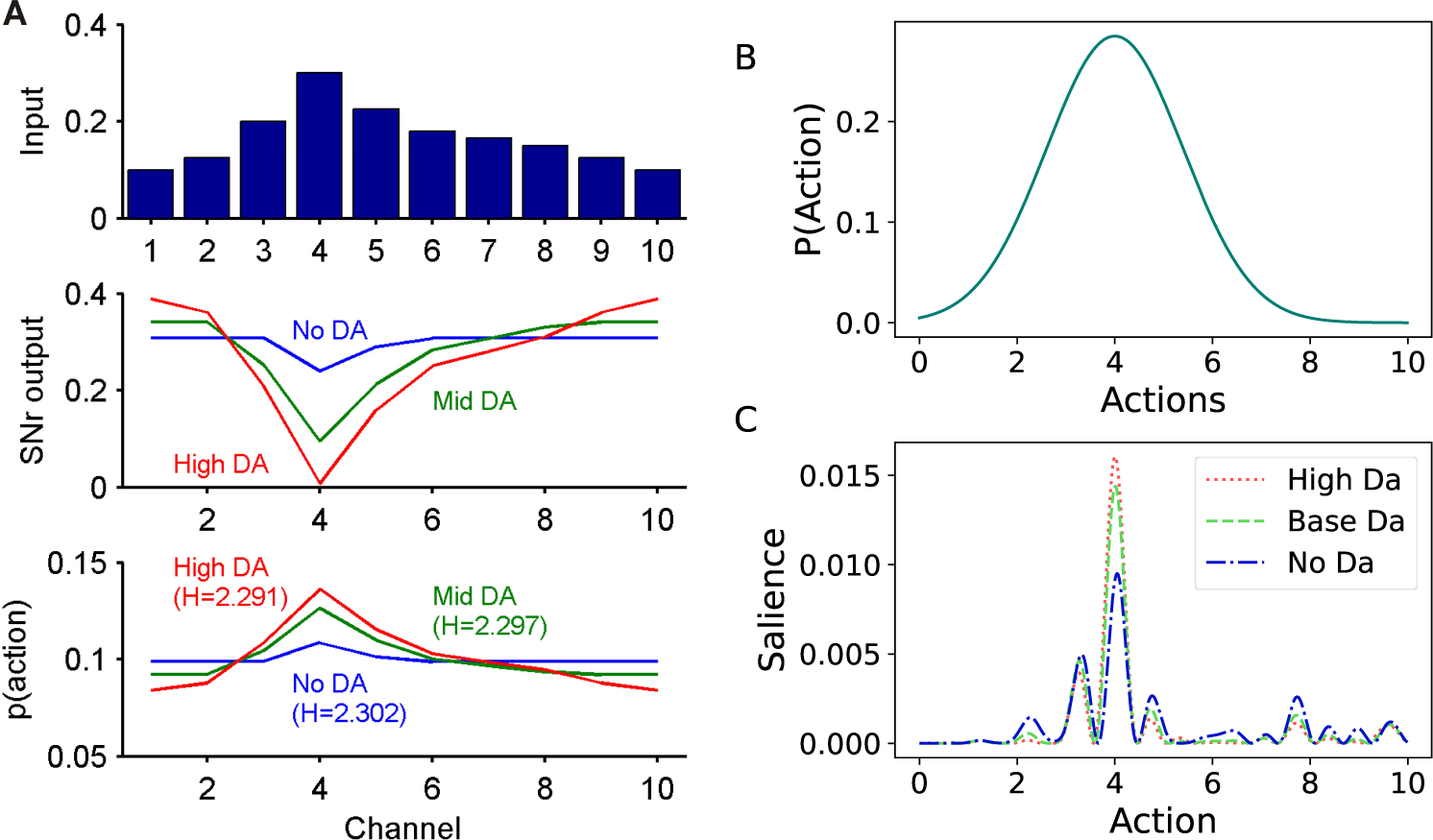
A. Figure 3A from Humphries et al. [60]. Shared via the CC-BY-NC3 license and provided at request by the authors. Top: distribution of inputs to the basal ganglia across 10 action channels. Middle: distribution of the output from the basal ganglia under 3 dopamine conditions (No = 0, Mid = 0.4, High = 0.8). Bottom: conversion from output salience to probability distribution function for action selection. Results illustrate that higher levels of dopamine corresponded to a probability distribution with a higher peak, indicating enhanced exploitation. H: entropy (in bits) of the PDF. B. Gaussian distribution over action space used as input to the basal ganglia system to test performance under different levels of dopamine. *µ* = 4.0*, σ* = 1.4. C. Cosine similarity (*S_c_*(*x, y*)) of basal ganglia model output under different dopamine conditions (No: 0, Mid: 0.4, High: 0.8). Results illustrate a replication of the findings of Humphries et al. [60] in that higher levels of tonic dopamine correspond to a higher peak, and lower tonic dopamine to a lower peak.

In our experiment, we used the same values for tonic dopamine (No = 0, Mid = 0.4 and High = 0.8) and set λ = 1.0 just as in [60]. We generated a similar input distribution, shown in Figure 9B in the form of a Gaussian distribution with parameters µ = 4.0, σ = 1.4. The results of this experiment are shown in Figure 9C. This plot illustrates that, as dopamine increases, the distribution being produced by the basal ganglia model is increasingly similar to an SSP encoding action ‘4’.

## 4 Discussion

The current work presents a novel model of action specification in the basal ganglia that provides an account of how continuously-valued action features, such as speed, might be represented in the brain. Furthermore, it offers an account of how the basal ganglia contributes to selection from these continuous action spaces, namely, by reducing the entropy of continuous salience distributions, thus reducing the range of actions available for a down-stream selection mechanism.

We leveraged VSAs, namely HRR, for encoding salience distributions over continuous action spaces. We do not claim that this is the only approach for encoding quasi-probability distributions over continuous variables. For example, this could be achieved by using methods such as log probability code [*e.g.*, 69], probability ratio codes [70], explicit probability codes [*e.g.*, 71, 72], convolutional probability codes [73–77], probabilistic population codes (PPCs) [78–82], or neural sampling methods [83–86].

The model was evaluated on 3 behavioural properties. The first and second properties concern our predictions on how the model should deal with uni- and bi-modal salience distributions. In the case of bi-modal distributions, it is important to acknowledge that not all instances of equally viable action options are resolved by simply selecting one of the modes. Rather, indecision can occur when an agent is presented with two equally or similarly salient options. This indecision can be momentary and manifest in the form of delayed responses or hesitations [87–89]. Evaluating the proposed model for its ability to capture these behaviours is beyond the scope of this paper.

The third property that we tested concerns the role of dopamine in modulating the explore-exploit trade-off. We were able to successfully replicate the results from [60], who tested the effect of modulating dopamine in the GPR model, by demonstrating the same relationship between dopamine and the tendency for exploration vs. exploitation in our model. Our results also represent a replication of experiments indicating the role of dopamine in the explore/exploit trade-off in human [90], and non-human animal behaviour [68, 91].

Overall, the proposed model shows promise for providing an account of previously under-explored basal ganglia behaviours, namely those concerning the control of *how* an action is executed. However, in its current state there are some limitations that are important to consider and which future work should address. We have already indicated some limitations, however these are not the only ones we feel it is important to discuss. First, we highlight that the model, as it is presented herein, does not contain a selection mechanism. That is, the output from the basal ganglia model is a distribution of salience over the available action space, not an unique action. As a result, we share the assumption presented in [60], wherein the authors propose that the output signal from the basal ganglia is used to inform a down-stream selection network, possibly one that receives converging inputs from both basal ganglia and cortex. In such a network, signals from basal ganglia could act as a dynamic threshold, such that the strength of cortical input necessary to activate the target nucleus is a function of the basal ganglia output. This assumption necessitates exploration into which downstream nuclei could perform the sampling function.

A second limitation of the current work is that we have so far only explored its behaviour in the context of continuous action spaces. Throughout this work we have proposed, however, that the basal ganglia may be organised in a mixed discrete-continuous fashion. High-level actions, such as turning, walking or grooming, may be represented as discrete actions using a localist encoding method, as indicated by neuroscientific research [12, 15, 40]. However, we propose that, within those localist channels, kinematic features of those actions, such as speed, might be represented as continuous, using a distributed representation. Testing this theory in the future necessitates that the model be adapted to accommodate mixed discrete-continuous distributions of salience. Happily, the VSA method employed is capable of such representations [92]. For example, one could use the VSA binding operation to bind continuous distributions of salience to vectors representing discrete action choices. Competition would then need to be resolved on two levels, first at the level of the discrete actions, and then for the continuous action features.

Third, we chose to implement a VSA embedding that produces a sinc kernel. An artifact of this choice is that the decoded salience distribution exhibits multiple local maxima. Whilst the chosen VSA embedding met the requirements of being a neurally instantiable distributed encoding method capable of representing continuous variables, our results reveal some potential limitations of this approach. Namely, as can be seen in Figure 8B, a consequence of the sinc kernel is that, whilst the network will select one mode over the other, there is a local maximum that peaks within the range of intermediate values. If we treat the action space as continuous, this means that, whilst there is a high probability of selecting from one of the two viable action ranges (i.e. 2.5-3.5), there is salience associated with some of the intermediate actions (2.0-2.5) as well as the high salience actions located under the left-hand salience peak (0.5-1.0). We can engineer the shape of the kernel that will describe similarity in the VSA space [93] by appropriately choosing the distribution of phases in our SSP. It is thereby possible to explore the viability and biological fidelity of a range of kernel shapes in accounting for basal ganglia behaviour. Whilst outside of the scope of the current work, we feel that this is an important avenue for future research.

Additionally, whilst not necessarily a limitation, further validation is required to establish how well this model accounts for behavioural and neuroscientific data. For example, one pressing next step is to investigate the model’s predictions about patterns of neuron activities. Our model relies on the existence of action-place cells in the striatum. The activities of these neurons are expected to replicate the findings of [7] and [6]. In these studies, the researchers found that neurons in the striatum exhibit unique speed-tuning curves. The action-place cells are constructed specifically to replicate this behaviour by being tuned to preferentially fire for specific points within the continuous action space. These same neurons in our model will also be active in representing similar/nearby points in the action space. Whilst this pattern of encoding is restricted to the striatum in the proposed model, it is unclear to these authors whether we should expect to find similar activation patterns in other regions of the basal ganglia.

The focus of this paper is on presenting the novel model as an account of the role of the basal ganglia in specifying action kinematics. Beyond the limitations highlighted above, there are a number of avenues for future work. When it comes to neurotransmitter effects, the current model addresses the role of dopamine but does not consider any other neurotransmitters at present. Recent work, however, has identified dissociable roles of dopamine and noradrenaline on the type of exploratory behaviour employed (directed vs. random) [94]. Future work can explore whether and what adaptations to the presented model are necessary to capture the different effects of these neurotransmitters. Additionally, the basal ganglia is known to be involved in reinforcement learning, and has often been model as the ‘actor’ portion of actor-critic models of learning [95]. We have ongoing work incorporating this novel model into reinforcement learning algorithms in order to control agents with continuous action spaces. This pursuit has the opportunity to provide insights into how biological agents learn to adapt the kinematics of their actions according to task demands. It will also demonstrate the utility of this approach for achieving continuous control for robotics and other agents in control tasks.

Collectively, the results reported in this paper demonstrate that the novel model of action specification in the basal ganglia offers promise in providing an account for previously under-explored basal ganglia behaviours. This work, therefore, provides an important and useful foundation for future exploration and validation experiments.

## Supplementary information

Not applicable

## Acknowledgements

Not applicable

